# Sucrose or sucrose and caffeine differentially impact memory and anxiety-like behaviours, and alter hippocampal parvalbumin and doublecortin

**DOI:** 10.1101/308684

**Authors:** Tanya J. Xu, Amy C. Reichelt

## Abstract

Caffeinated sugar-sweetened “energy” drinks are a subset of soft drinks that are popular among young people worldwide. High sucrose diets impair cognition and alter aspects of emotional behaviour in rats, however, little is known about sucrose combined with caffeine. Rats were allocated to 2h/day 10% sucrose (Suc), 10% sucrose plus 0.04% caffeine (CafSuc) or control (water) conditions. The addition of caffeine to sucrose appeared to increase the rewarding aspect of sucrose, as the CafSuc group consumed more solution than the Suc group. After 14 days of intermittent Suc or CafSuc access, anxiety was assessed in the elevated plus maze (EPM) prior to their daily solution access, whereby CafSuc and Suc rats spent more time in the closed arms, indicative of increased anxiety. Following daily solution access, CafSuc, but not Suc, rats showed reduced anxiety-like behaviour in the open-field. Control and CafSuc rats displayed intact place and long-term object memory, while Suc showed impaired memory performance. Sucrose reduced parvalbumin immunoreactivity in the hippocampus, but no differences were observed between Control and CafSuc conditions. Parvalbumin reactivity in the basolateral amygdala did not differ between conditions. Reduced doublecortin immunoreactivity in the dentate gyrus relative to controls was seen in the CafSuc, but not Suc, treatment condition. These findings indicate that the addition of caffeine to sucrose attenuates cognitive deficits. However, the addition of caffeine to sucrose evokes anxiety-like responses under certain testing conditions, suggesting that frequent consumption of caffeinated energy drinks may promote emotional alterations and brain changes compared to standard soft drinks.

## 1. Introduction

Energy drinks, high in both sugar and caffeine, are consumed frequently by young people (Zucconi et al., 2013). Overconsumption of sugar-sweetened beverages, including soft-drinks, is associated with poor physical health outcomes such as obesity and type-2 diabetes (Te Morenga et al., 2014). The addition of caffeine to sugar-sweetened beverages in the form of “energy drinks” may further exacerbate physical and mental health problems (Koivusilta et al., 2016). Caffeine is a cognitive stimulant and can enhance the rewarding properties of soft drinks (Costa et al., 2014). Caffeine acts as an antagonist at adenosine receptors and potentiates the release of neurotransmitters involved in reward, mood, and cognition (Fredholm et al., 1999). Regular caffeine intake persistently antagonises A_1_ and A_2a_ adenosine receptors, upregulating adenosine A_1_ receptors in the amygdala and hippocampus, which may underpin changes in emotion and cognition (Shi et al., 1993).

Sucrose and caffeine consumption separately impact cognitive performance in rodents (e.g., Ardais et al., 2016; Reichelt et al., 2016). Daily access to sucrose, either intermittently or continuously disrupts place recognition memory, but not object recognition memory with short (5 min) retention intervals, indicative of impaired hippocampal, but no perirhinal cortex, function (Abbott et al., 2016; Beilharz et al., 2016; Kendig et al., 2013). Chronic sucrose intake impaired hippocampal-mediated long-term object recognition memory in adolescent rats when retention intervals were extended to 1h (Jurdak and Kanarek, 2009). Moreover, intermittent (2h) daily access to 10% sucrose reduced hippocampal immunoreactivity in parvalbumin (PV)-expressing GABAergic interneurons (Reichelt et al., 2015) and doublecortin in the dentate gyrus (DG) (Reichelt et al., 2016), indicating that sucrose impacts on aspects of inhibitory neurotransmission and neuroproliferation in the hippocampus. In contrast, caffeine consumption has been shown to improve memory performance in rats (Ardais et al., 2014; Costa et al., 2008). Intermittent (12h / day) access to 0.04% or 0.08% caffeine for 20 days improved perirhinal-mediated object recognition memory (Ardais et al., 2014) and 90 days of caffeine supplemented chow selectively enhanced long-term object recognition memory (Abreu et al., 2011). However, little is known how the combination of caffeine and sucrose impacts on behaviour, and what neurobiological changes might underpin these effects.

Anxiety has been linked to frequent energy drink intake in young people (Stasio et al., 2011; Trapp et al., 2014). Separately, high sucrose diets and caffeine-supplemented diets in rats evoke anxiety-like behaviours (e.g., Avena et al., 2008; O’Neill et al., 2016). Ardais et al. (2014) reported that adolescent male rats with continuous access to caffeine for 25 days displayed increased anxiety-like behaviour in the elevated plus maze (EPM). Adolescent (but not adult) male rats with chronic 0.03% caffeine intake for 28 days showed heightened anxiety-like behaviour in the EPM and open-field and increased amygdala and hypothalamus cortisol expression following 7 days caffeine abstinence, mimicking aspects of caffeine dependency (O’Neill et al., 2016).

Despite the generally cognitive enhancing properties of caffeine, sugar-sweetened energy drinks have been linked to both positive (Wesnes et al., 2017) and negative (Van Batenburg-Eddes et al., 2014) cognitive outcomes. A recent study suggested that the addition of caffeine to sucrose solution may protect against sucrose-induced deficits in hippocampal-mediated memory, as mice with 10% sucrose + 0.03% caffeine access for 30 days showed intact place memory in the Y-Maze (Nieradko-Iwanicka et al., 2016).

In this study, we sought to determine the impact of daily intermittent sucrose and sucrose plus caffeine on anxiety and memory in young male rats. As previous studies have linked reduced parvalbumin and doublecortin immunoreactivity to intermittent sucrose access induced cognitive deficits (Reichelt et al., 2015; Reichelt et al., 2016) we examined the effect of sucrose and sucrose plus caffeine examined these markers in the hippocampus and basolateral amygdala (BLA). Parvalbumin-expression in the BLA is linked to anxiety-like behaviour (Urakawa et al., 2013), as such we also measured parvalbumin-immunoreactivity across treatment groups in this region.

## 2. Method

### 2.1. Subjects

Three-week-old male Sprague-Dawley rats (*N* = 36) were supplied by the Animal Resources Center, Western Australia. Rats were group-housed, 4 per plastic cage (26×40×60 cm), in a 21±2 °C colony room on a 12h light/dark cycle (lights on at 7:00 am). Rats were weight-matched across diet groups and allocated to Control (*N* = 16), Sucrose (Suc; *N* = 8) or Sucrose plus caffeine (CafSuc; *N* = 12) conditions. Body weight and 24h chow intake (converted into kJ) were measured twice weekly. Experimental procedures were approved by the RMIT University Animal Ethics Committee.

### 2.2. Diet conditions

Starting at postnatal (P) day 28, Suc and CafSuc treatment groups received 2h daily access (9:00-11:00am) to a bottle of 10% sucrose (w/v) or 10% sucrose + 0.04% caffeine (w/v) solution in their home-cages across a 28 day period (P28-P55) that encompassed adolescence (Spear, 2000). Sucrose and caffeine concentrations and the caloric densities (1.7 kJ/ml) of the solutions were comparable to commercial sugar-sweetened soft drinks and energy drinks (e.g. Red Bull, Monster, V). A caffeine alone group was not used in this study due to the bitter nature of 0.04% caffeine, which was found to not be readily consumed by rats over the 2h access period. It was deemed that other routes of caffeine administration (oral gavage, intraperitoneal injections) would not be comparable to the 2h limited access protocol in the Suc and CafSuc conditions due to potential stress or altered pharmacokinetics. Consumption of the solutions was recorded daily by weighing bottles before and after access. All rats had *ad-libitum* access to standard laboratory chow (11 kJ/g) and water.

### 2.3. Behavioural Procedures

Figure 1 outlines the experimental timeline. Behavioural testing commenced after the rats had been exposed to Suc and CafSuc solutions for 2h/day for 14 days. Each rat was tested on the elevated plus maze (EPM), open-field, novel place recognition (NPR) and short-and long-term novel object recognition (NOR) tasks across experimental days 14-28, between 9:00am-2:00pm. Performance in the EPM and open-field relies upon novelty of exploring an unfamiliar environment to quantify anxiety-like behaviours. As such, Suc and CafSuc rats were tested in the EPM prior to receiving their daily solution access (~24h solution-deprived). Subsequent behaviours, including the open-field, were tested after solution access. Arenas were cleaned with 70% ethanol between test phases to remove odour cues and illuminated at 20 Lux. A video camera positioned above each apparatus recorded behaviour.

#### Elevated plus maze

Rats underwent testing in once in the EPM as a measure of exploration in a novel environment (Walf and Frye, 2007). The maze comprised of four arms (50×10cm) joined by a centre platform (10×10cm) to form a plus shape elevated 50cm off the ground. Two opposing arms were enclosed by 40cm tall walls and the other two had no walls. Each rat, placed in the centre, could explore all arms for 5 min. Time spent in closed arms (Time_closed_), centre (Time_centre_) and open arms (Time_open_) were scored using ODLog (Macropod Software) and converted into a percentage of the total time in the EPM, %Time_closed_ quantified anxiety. Further ethological parameters were assessed including head dips (downward movement of rodents’ head toward the floor from the open arms), rearing (vertical standing on two hind legs), freezing behaviour (when the rodent is motionless and ears pointed forward), grooming, and risk assessment defined by time in a stretched posture into the open arm with hind legs in the closed arm.

#### Open-field

Rats underwent testing in once in the open field as a measure of exploration in a novel environment. The open-field was a square arena (50×50×60cm) constructed from black Perspex. Each rat could explore the open-field for 5 min. Total distance travelled (Distance_total_) and percentage of time spent in arena border (%Time_border_) and centre (%Time_centre_) were scored using SMART video-tracking system (Panlab, Harvard Systems, USA). Distance_total_ quantified locomotor activity, and %Time_border_ quantified anxiety.

#### Novel Place Recognition (NPR) and Novel Object Recognition (NOR) tasks

Commercial objects (e.g., bottles) of differing heights, widths, and colour were used as stimuli within the square open-field arena. Figure 1B and C illustrates the NPR and NOR tasks. Each rat was placed in the arena and could explore two identical objects for 5 min (sample phase). Rats were then removed for a 5 min (NPR and NOR)(NPR and NOR) or 24h (long-term NOR) retention interval. Rats were returned to the arena (3 min, test phase) and could explore a familiar and novel located sample object (NPR), or a familiar and novel object (NOR). Object exploration times were scored using ODLog to calculate exploration ratios (eRs), eR = Time_novel_ / (Time_novel_ + Time_familiar_). An eR greater than 0.5 indicated preference for the novel location/object and intact memory. An eR ≤ 0.5 indicated an equal preference for novel and familiar locations/objects, or preference for the familiar object, and therefore impaired memory.

**Figure 1.**
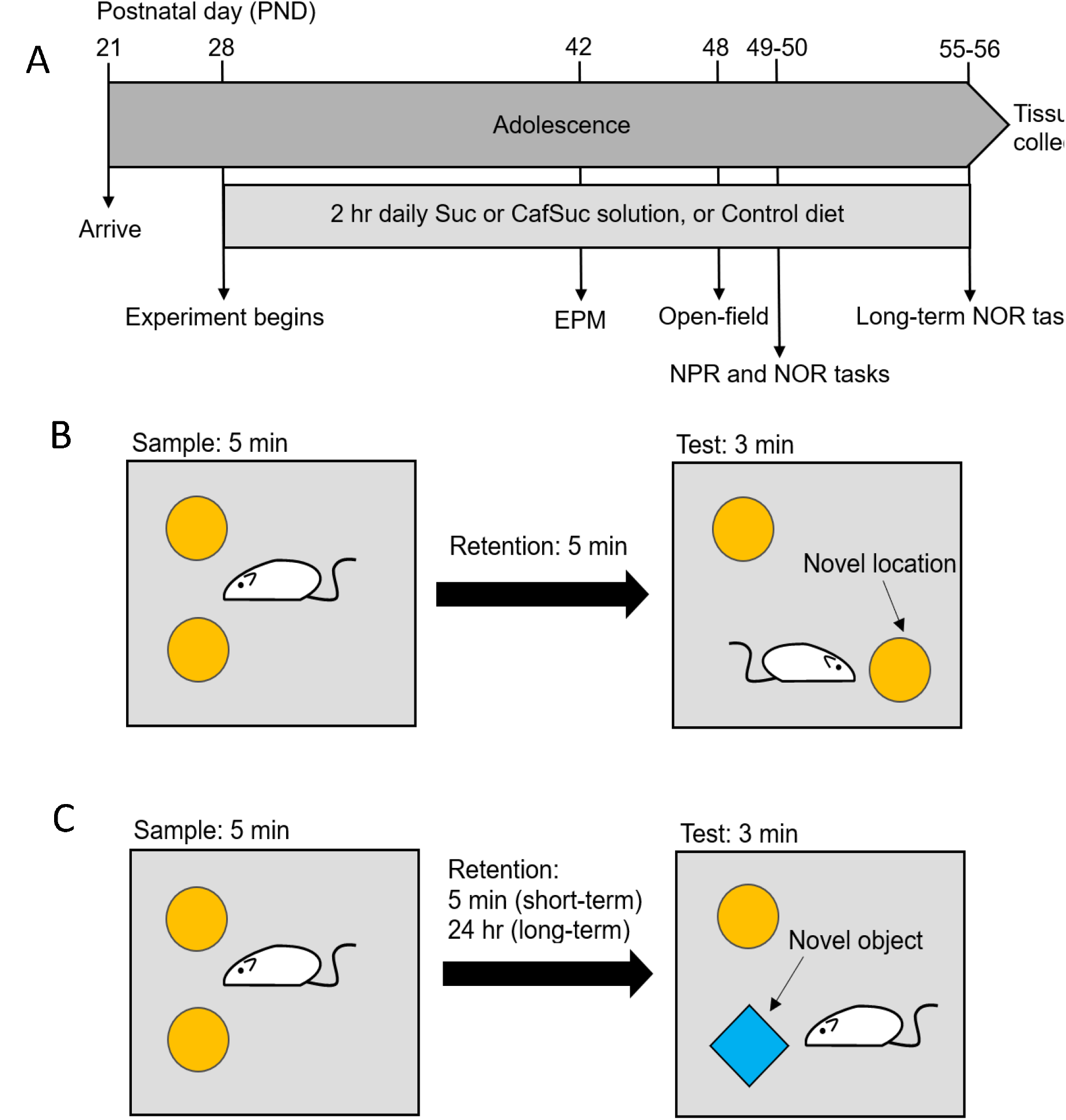
(A) Timeline of experimental study. Suc = 10% sucrose (w/v); CafSuc = 10% sucrose + 0.04% caffeine (w/v); EPM = elevated plus maze; NPR = novel place recognition; NOR = novel object recognition. Schematic representation of the (B) Novel place recognition (NPR) task and (C) Novel object recognition (NOR) task (short-and long-term).

### 2.4. Immunohistochemistry and Immunofluorescence

Following behavioural testing, rats were deeply anesthetized with sodium pentobarbital (100 mg/kg ip) and perfused transcardially with 0.9% saline containing heparin (5000 IU/ml), followed by 4% paraformaldehyde (PFA) in 0.1 M phosphate buffer (PB), pH 7.4. Brains were postfixed for 24h in 4% PFA and then cryoprotected in PB plus 20% sucrose for 48 hrs. Brains were sliced into 40 μm coronal sections on a cryostat into a series of 4 slices, thus slices from each series were separated by 160 μm. Sections containing the dorsal hippocampus and BLA were stained for parvalbumin immunoreactivity or doublecortin immunofluorescence as described below.

#### Parvalbumin immunohistochemistry

Sections were washed in 0.1M phosphate buffered saline solution (PBS) and blocked with 4% normal horse serum (NHS) diluted in 0.1 M PBS containing 0.1% Triton X-100 (PBS-T), then incubated at 4°C with a mouse monoclonal antibody against parvalbumin (Millipore, MAB1572) at a dilution of 1:5000 in PBS-T plus 2% NHS for 48hr. Washed sections were blocked for endogenous peroxidase activity with hydrogen peroxide (0.1%) and incubated with a biotinylated secondary antibody (horse anti-mouse, Vector Laboratories, dilution 1:500 in 2% NHS). Sections were incubated with Streptavidin-Horseradish Peroxidase (1:500, in PBS) and visualized using 3’,3’-diaminobenzidine (DAB) intensified with nickel chloride. Sections were mounted onto 4% gelatin-coated slides, dehydrated in ethanol, cleared in histolene, and cover-slipped with DPX (Sigma). Parvalbumin immunoreactive neurons were imaged using an automated slide-scanner (Olympus VS120-S5, 10× objective lens, Olympus VS-ASW software). Total counts of parvalbumin immunoreactive cells (PV+ cells) were made from 3-4 sections per animal from the left hemisphere using ImageJ (v1.46; http://imagejnih.gov/ij). Counts were made from the dorsal hippocampus (between bregma - 2.6mm and −3.8mm) separated into the CA1, CA3 and DG regions, and the BLA (lateral and basal nuclei, between bregma −2.2 mm and −3.4 mm). Regions were delineated using clearly visible landmarks and predefined boundaries according to a rat brain atlas (Paxinos and Watson, 2013).

#### Doublecortin immunofluorescenc

Sections were washed in 0.1 M PBS and blocked with 4% normal rabbit serum (NRS) diluted in 0.1 M PBS-T, then incubated at 4°C with a goat monoclonal antibody against doublecortin (Santa Cruz, C10, 1:500) PBS-T + 2% NRS for 48h. Sections were then washed in PBS and incubated with a fluorescent secondary antibody (AlexaFluor rabbit anti-goat 488, dilution 1:2000 in PBS-T + 2% NRS) for 2h at room temperature. Sections were mounted onto 4% gelatin-coated slides and cover-slipped with Vectorshield + DAPI. Doublecortin-expressing neurons were imaged using a stage-controlled fluorescent microscope (Olympus BX61, 20× objective lens, Nikon NIS-Elements software). Doublecortin was quantified in ImageJ by calculating the percentage of a 1mm^2^ area (%Area_DCX_) within the granule cell layer of the DG in coronal hippocampal slices between bregma −2.8mm and −3.6mm (see Fig 7B). For this analysis, the threshold for positive staining exceeding −10 to −255 (black) and minimum 1μm area criterion was applied by densitometric scanning of the targets. The magnitude of doublecortin immunofluorescence was quantified as the average area in the positive threshold from 4 sections of the left hemisphere.

### 2.5. Statistical Analysis

Statistical comparisons were conducted in IBM SPSS v24, **α** = .05. Body-weight, consumption and sample phase object exploration (NPR/NOR tasks) were analysed using mixed-design analysis of variances (ANOVAs) with within-subjects (time or object) and a between-subjects factor of diet (Control/Suc/CafSuc). Simple main effects analyses (Bonferonni adjusted) were used to explore significant interactions. Baseline body-weight and data from the EPM (%Time_closed_, %Time_centre_, %Time_open_), open-field (Distance_total_, %Time_border_, %Time_centre_), Exploration ratios from memory tasks, parvalbumin immunopositive cells (#PV+ cells) and immunofluorescence (%Area_DCX_) were analysed using one-way ANOVAs for the effects of diet. A Greenhouse-Geisser correction was applied upon sphericity violation. Welch’s *F* was reported when homogeneity of variances was violated. *Post hoc* Tukey-Kramer tests explored significant main effects. Data were expressed as Mean ±*SEM*.

## 3. Results

### 3.1. Body-Weight

Figure 2A shows the mean body-weight of rats across groups throughout the experiment. Groups showed similar baseline body-weights (main effect of diet *F* < 1). Mean body weights of rats increased at a similar rate between groups, demonstrated by a significant main effect of time, *F*(1.18, 38.9) = 1227.1, *p* < .001, but no significant main effect diet, *F* < 1, or significant time×diet interaction, *F* < 1.

**Figure 2.**
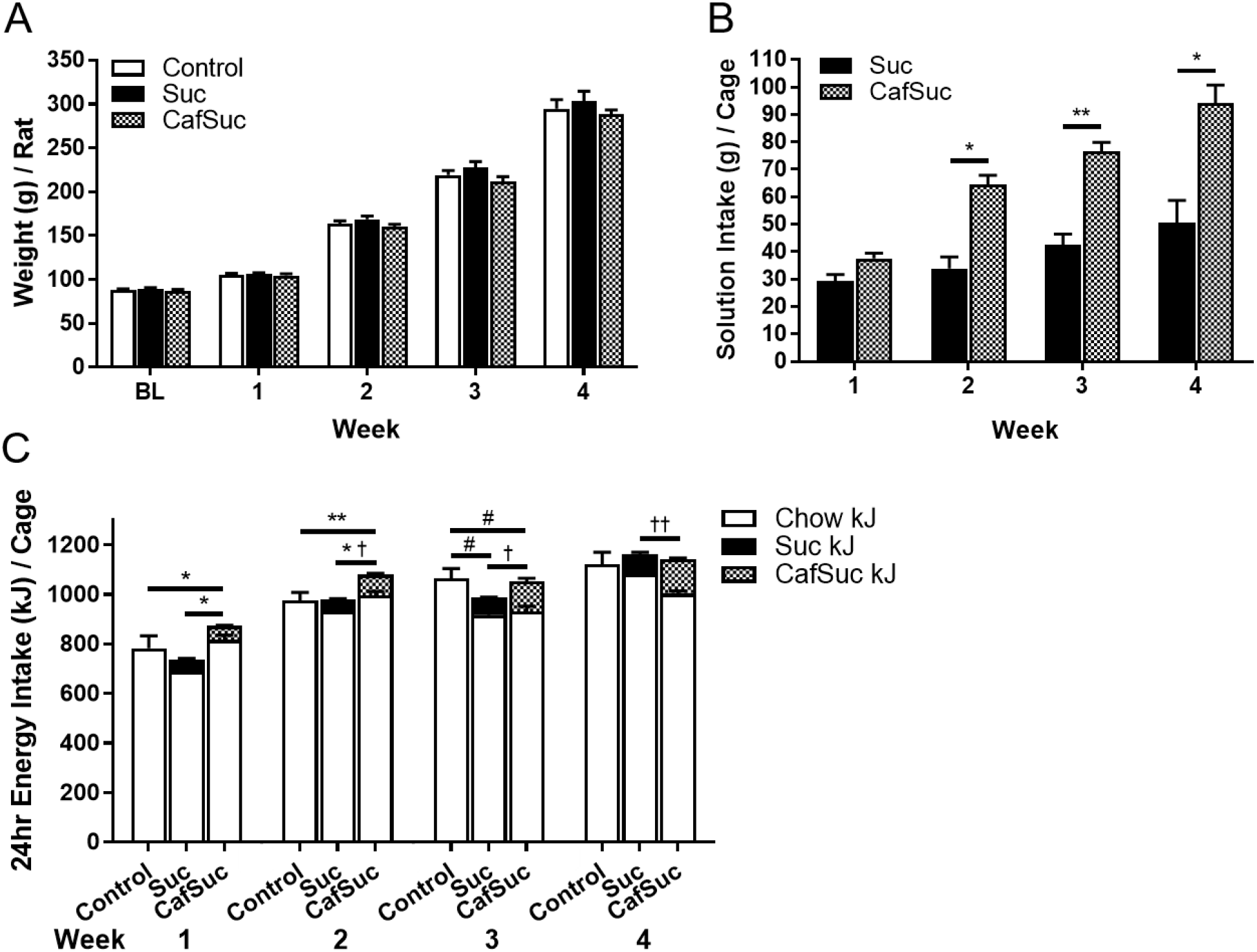
Physiological data across the four weeks of the experiment. (A) Mean body-weights (g) of rats in Control (*N* = 16), Suc (*N* = 8) and CafSuc (*N* = 12) groups. (B) Mean amount of sucrose or sucrose plus caffeine solution (g) consumed per cage during the 2h daily access period across Suc and CafSuc treatment groups. (C) Mean 24h energy intake (kJ) per cage across Control (*ad-libitum* chow), Suc (*ad-libitum* chow with intermittent sucrose) and CafSuc (*ad-libitum* chow with intermittent sucrose plus caffeine) groups. BL = baseline; Suc = Sucrose treatment group; CafSuc = Sucrose plus caffeine treatment group; Suc kJ = sucrose solution kJ; CafSuc kJ = sucrose plus caffeine solution kJ. Solution intake (g): \**p* < .05, \*\**p* < .01. Significant difference in total energy intake: \**p* < .05, \*\**p* < .01. Significant differences between Control and Suc, and Control and CafSuc chow energy intake: #*p* < .05. Suc and CafSuc solution energy intake: †*P* < .05, ††*P* < .01. Error bars represent ±*SEM*.

### 3.2. Consumption

Figure 2B illustrates daily consumption of solutions (Suc/CafSuc) in 2h per cage throughout the experiment. Solution consumption differed between Suc and CafSuc groups, indicated by a significant time×diet interaction, *F*(3,9) = 16.7, *p* < .001. CafSuc solution intake exceeded that of Suc from week 2 onwards: week 2 (*F*(1, 3) = 31.5, *p* = .01); week 3 (*F*(1, 3) = 45.7, *p* = .007); week 4 (*F*(1, 3) = 16.7, *p* = .03).

Total 24h energy intake increased at a similar rate between groups (see Figure 2C), indicated by a significant main effect of time, *F*(1.28, 7.66) = 103.9, *p* < .001, but no significant main effect of diet, *F*(2, 6) = 1.23, *p* = .36, or time×diet interaction, *F*(2.55, 7.66) = 2.84, *p* = .11. However, chow 24h energy intake differed between groups across time, indicated by a significant time×diet interaction, *F*(6, 18) = 5.04, *p* < .001. Control chow energy intake exceeded that of Suc (*p* = .03) and CafSuc (*p* = .03) in week 3. Daily 2h solution energy intake differed between Suc and CafSuc groups across time, indicated by a significant time×diet interaction, *F*(3, 9) = 9.1, *p*= .004. CafSuc solution energy intake exceeded that of Suc from week 2 onwards: week 2 (*F*(1, 3) = 18.6, *p* = .02); week 3 (*F*(1, 3) = 11.8, *p* = .04); week 4 (*F*(1, 3) = 48.5, *p* = .006).

### 3.3. Elevated Plus Maze

Groups differed in %Time_closed_ as shown in Figure 3A, demonstrated by a significant main effect of diet, *F*(2, 33) = 8.5, *p*= .001. Suc and CafSuc rats showed increased anxiety-like behaviour, spending significantly greater %Time_closed_ than Controls (Suc *p* = .004; CafSuc *p*= .006), but comparable %Time_closed_ to each other (*p* = 1.00). Figure 3B shows that groups also differed in %Timecentre (significant main effect of diet, *F*(2,33) = 11.0, *p* < .001). Suc and CafSuc rats spent significantly less %Time_centre_ than Controls (Suc *p* = .001; CafSuc *p* = .002), but comparable %Time_centre_ to each other (*p* = .73). There were no group differences in %Time_open_, *F*(2, 20.28) = 1.7, *p* = .21 (see Figure 3C).

Groups differed in time spent head dipping, engaged in risk assessment and rearing but not in grooming and freezing behaviour in the EPM as shown in Table 1.

**Table 1.** Effect of diet treatments on the duration (s) Elevated Plus Maze behaviours - grooming, head dipping, risk assessment, rearing and freezing. Suc = Sucrose group. CafSuc = Caffeine + Sucrose group. \**p* < .05. \*\**p* < .01.

| Behaviour | Group | Duration (s) | | $F(df)$ | $p$ |
| --- | --- | --- | --- | --- | --- |
|  |  | Mean | SEM |  |  |
| Grooming | Control | 11.24 | 6.20 | 0.65(2, 33) | .529 |
|  | Suc | 8.94 | 2.63 |  |  |
|  | CafSuc | 3.82 | 0.96 |  |  |
| Head Dipping | Control | 4.97 | 1.24 | 5.84(2, 33)** | .007 |
|  | Suc | 2.99 | 1.06 |  |  |
|  | CafSuc | 3.09 | 0.89 |  |  |
| Risk Assessment | Control | 15.27 | 3.82 | 5.07(2, 33)* | .012 |
|  | Suc | 6.53 | 2.31 |  |  |
|  | CafSuc | 11.81 | 3.41 |  |  |
| Rearing | Control | 15.29 | 1.30 | 8.91(2, 33)** | .001 |
|  | Suc | 12.35 | 1.69 |  |  |
|  | CafSuc | 23.09 | 2.24 |  |  |
| Freezing | Control | 1.47 | 0.37 | 1.52(2, 19.27)<sup>a</sup> | .244 |
|  | Suc | 0.49 | 0.18 |  |  |
|  | CafSuc | 6.10 | 1.76 |  |  |

<sup>a</sup>Welch's $F$ used due to violation of homogeneity of variances assumption.

Table 2 shows that the Control group spent significantly more time engaged in head dipping than CafSuc and Suc groups. Additionally, Control and CafSuc rats spent significantly greater time engaged in risk assessment behaviour than Suc rats. Finally, Control and Suc rats spent significantly less time rearing than CafSuc rats.

**Table 2.** Post hoc comparisons between groups for head dipping, risk assessment and rearing behaviour in the Elevated Plus Maze. \**p* < .05. \*\**p* < .01. Suc = Sucrose group. CafSuc = Caffeine + Sucrose group.

| Behaviour | Group 1 | Group 2 | Difference<sup>a</sup> |  | *p* |
| --- | --- | --- | --- | --- | --- |
|  |  |  | *Mean* | *SEM* |  |
| Head Dipping | Control | Suc | 4.44* | 1.75 | .041 |
|  | Control | CafSuc | 4.74* | 1.54 | .011 |
|  | Suc | CafSuc | 0.30 | 1.84 | 1.000 |
| Risk Assessment | Control | Suc | 14.93* | 5.50 | .028 |
|  | Control | CafSuc | -2.51 | 4.85 | 1.000 |
|  | Suc | CafSuc | -17.45* | 5.80 | .014 |
| Rearing | Control | Suc | 2.84 | 2.64 | .511 |
|  | Control | CafSuc | -7.79** | 2.33 | .006 |
|  | Suc | CafSuc | -10.73** | 2.78 | .001 |

<sup>a</sup>Group 1 – Group 2

### 3.4. Open-Field

Figures 3D show that groups differed in %Time_border_ (significant main effect of diet, *F*(2, 33) = 11.5, *p* < .001). CafSuc treatment rats spent significantly less %Time_border_ than Control (*p*< .001) and Suc (*p* = .001) treatment groups, indicative of reduced anxiety-like behaviour. However, groups did not differ in distance travelled (see Fig 3E), demonstrated by no significant main effect of diet, *F*(2, 33) = 1.6, *p* = .21, indicating similar locomotor activity.

**Figure 3.**
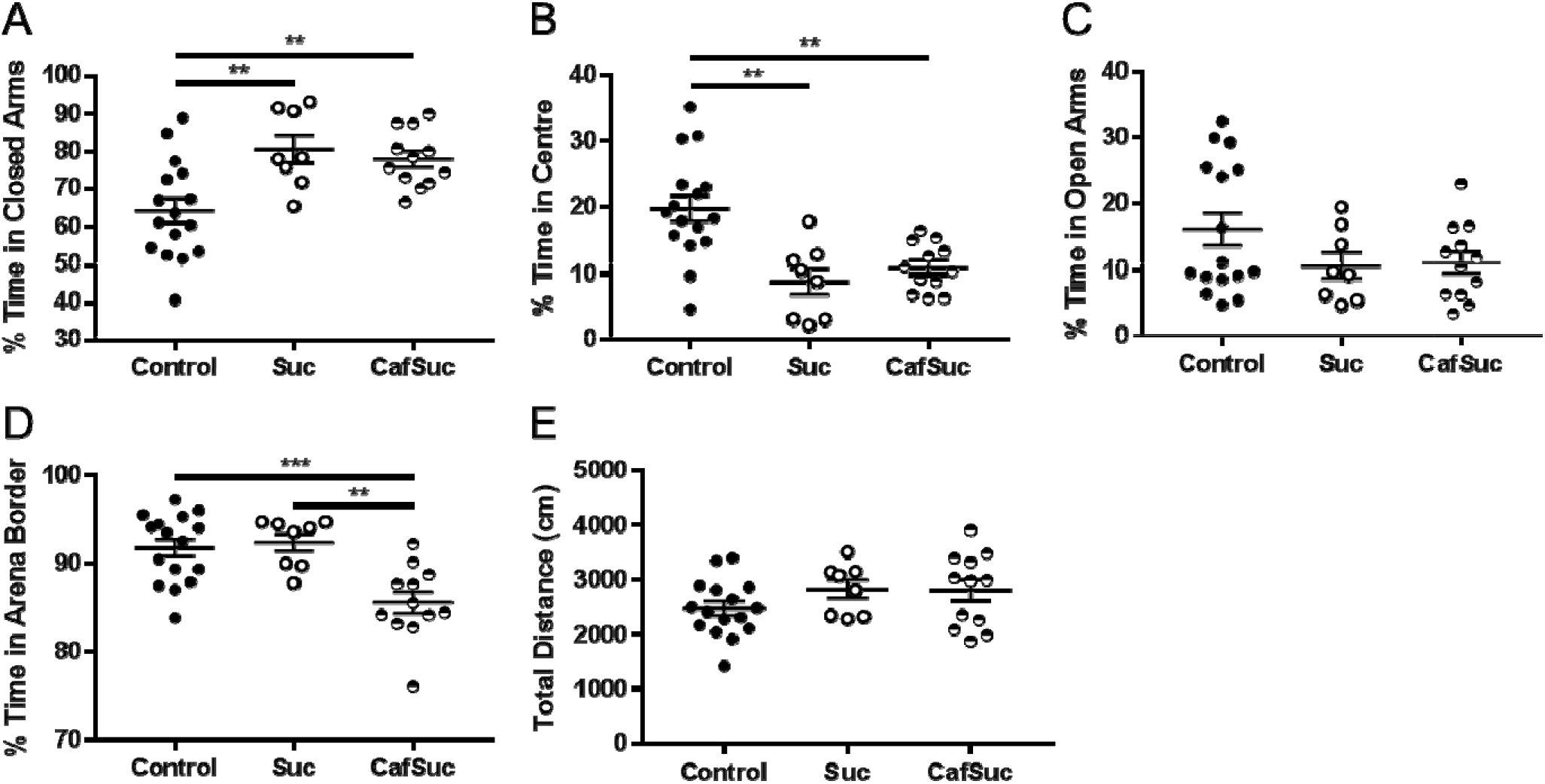
(A) Percentage of time spent in closed arms (%Time_closed_), (B) centre (%Time_centre_) and (C) open arms (%Time_open_) of the elevated plus maze (EPM) across Control (*N* = 16), Suc (*N* = 8) and CafSuc (*N* = 12) groups. (D) Percentage of time spent in the border (%Time_border_) and (E) total distance travelled (cm) in the open-field arena across Control (*N*= 16), Suc (*N*= 8) and CafSuc (*N* = 12) groups. \*\**p* < .01. \*\*\**p* < .001. Data shows group means and error bars represent ±*SEM*.

### 3.5. Novel place recognition and novel object recognition performance

Rats were excluded from analyses if they explored each object for less than 1 s during sample/test phases, resulting in *N* = 33 (novel place recognition), *N* = 34 (short-term novel object recognition) and *N* = 34 (long-term novel object recognition).

#### Sample phases

Rats explored the sample objects equally [no significant main effects of object: NPR *F*(2, 31) = 2.9, *p* = .10; short-term NOR *F*< 1; long-term NOR *F*(2, 31) = 2.6, *p*= .115; or significant time×diet interaction, NPR *F* < 1; short-term NOR, *F* < 1; long-term NOR, *F* < 1]. However, group differences were observed in sample object exploration in the long-term NOR [Mean (±SEM) Control = 13.0 (1.6), Suc 24.1 (2.2), CafSuc = 16.8 (1.8); significant main effect of diet *F*(2,31) = 8.6, *p* = .001] but not for the NPR [Mean (±SEM) Control = 13.1 (2.1), Suc = 19.9 (2.7), CafSuc = 16.1 (2.2); no significant main effect of diet, *F*(2,31) = 2.1, *p* = .14] or short-term NOR [Mean (±SEM) Control = 11.2 (1.8), Suc = 16.4 (2.7), CafSuc = 16.1 (2.1); no significant main effect of diet, *F*(2,31) = 2.1, *p* = .14].

### Test phases

#### Novel Place Recognition

Groups differed in place eRs as shown in Figure 4A, indicated by a significant main effect of diet, *F*(2, 30) = 5.3, *p* = .01. The Suc group showed an equal preference for novel and familiar located objects, indicative of impaired spatial memory. Control and CafSuc groups demonstrated intact place memory (eRs > 0.5), with significantly greater eRs compared to the Suc treatment group (Control *p* = .032, CafSuc *p* = .011), but comparable eRs to each other (*p* = .80).

#### Short-term Novel Object Recognition

All conditions displayed intact short-term novel object recognition. No significant differences were observed between treatment groups as shown in Figure 4B (*F* < 1). All groups had eRs greater than 0.5, indicating a preference for the novel object.

#### Long-term Novel Object Recognition

Groups differed in long-term object memory as shown in Figure 4C, indicated by a significant main effect of diet, *F*(2, 31) = 4.97, *p* = .01. CafSuc treatment groups demonstrated intact long-term object memory (eR > 0.5), exhibiting greater eRs than Suc treatment groups (*p* = .010) and comparable eRs to Controls (*p* = .16). The Suc treatment group did not differ significantly to Controls (*p* = .26), however had a mean eR of 0.5, indicating equal preference for novel and familiar objects and impaired long-term object memory.

**Figure 4.**
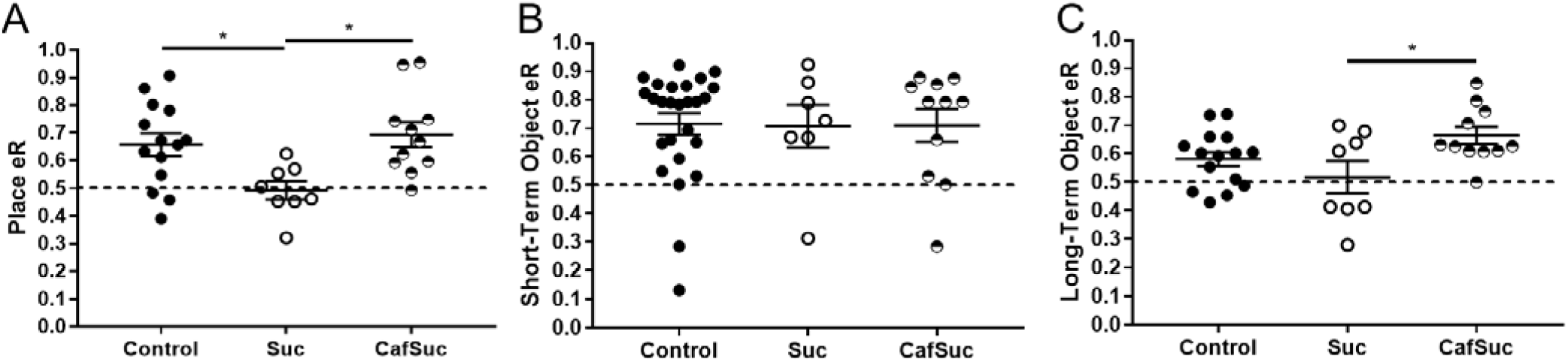
Test phase object / place exploration ratios (eR) in the (A) novel place recognition task, (B) short-term novel object recognition task and (C) long-term novel object recognition task across Control, Suc and CafSuc treatment groups. Suc = Sucrose group; CafSuc = Sucrose plus caffeine group; eR = exploration ratio: Time_novel_ / (Time_novel_ + Time_familiar_); Dotted line = 0.5 (equal preference between novel vs. familiar locations / objects). Data show means and error bars represent +*SEM*. \**p* < .05.

### 3.6. Parvalbumin immunoreactivity

#### 3.6.1. Hippocampus

Groups significantly differed in counts of overall dorsal hippocampal PV+ cells (see Figure 5A; significant main effect of diet, Welch’s *F*(2, 33) = 6.5, *p* = .004) with the Suc treatment group displaying significantly fewer overall hippocampal #PV+ cells than Control (*p* = .005) and CafSuc (*p* = .001) treatment groups. Groups differed in counts of PV+ cells in hippocampal subfields (see Figure 5C) CA1 (significant main effect of diet, Welch’s *F*(2, 33) = 20.3, *p* = .03) and CA3 (significant main effect of diet, Welch’s *F*(2, 33) = 20.2, *p* < .001) but not DG (no significant main effect of diet, *F*(2, 33) = 2.3, *p* = .12). Controls had significantly more CA1 #PV+ cells than Suc (*p* = .02) but comparable CA1 #PV+ cells to CafSuc (*p* = .82) rats. Control and CafSuc treatment groups displayed significantly more CA3 #PV+ cells than Suc (Control *p* < .001; CafSuc *p* = .004) and comparable CA3 #PV+ cells to each other (*p* = .58).

**Figure 5.**
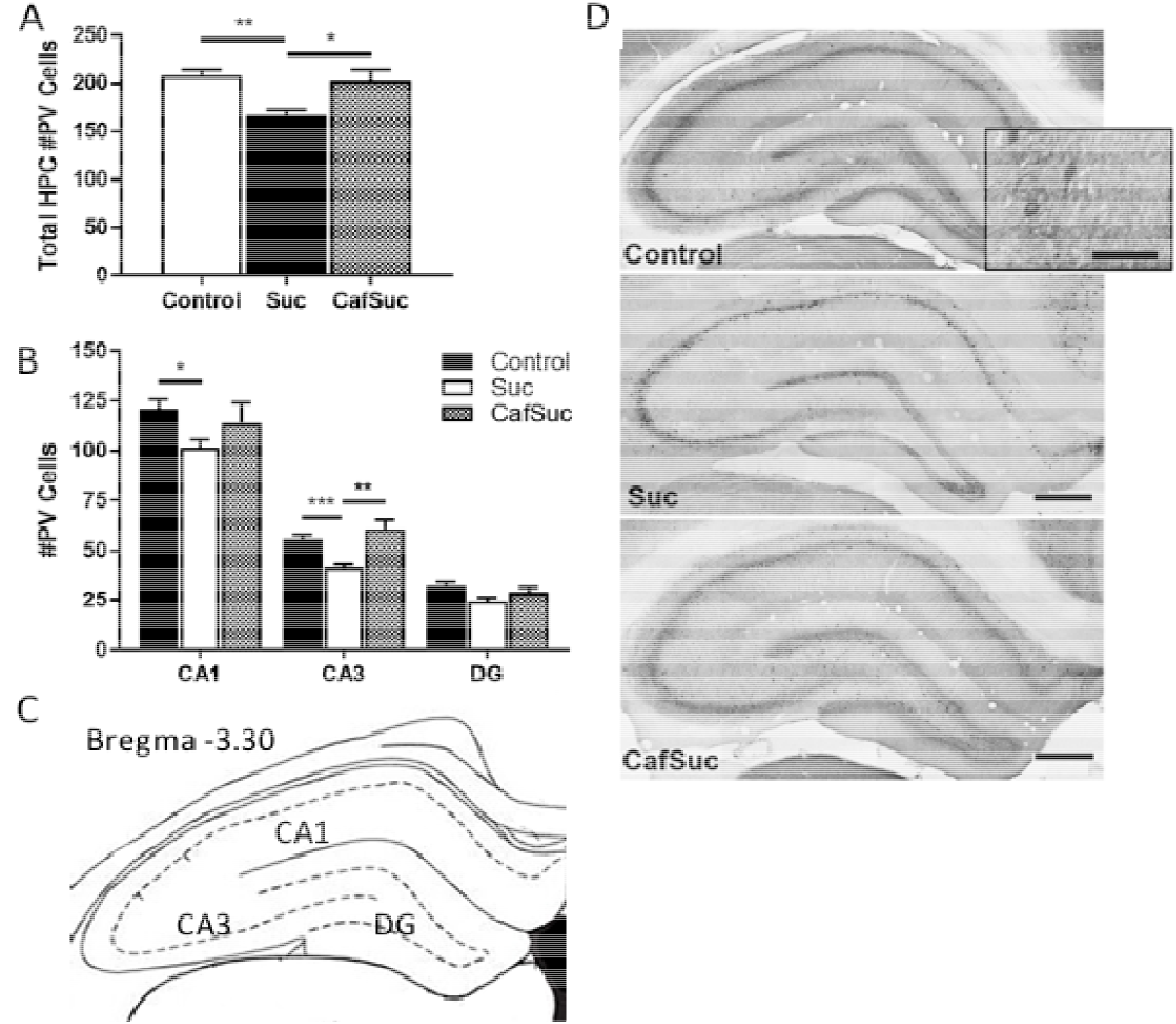
Dorsal hippocampal parvalbumin immunoreactivity in Control, Suc and CafSuc treatment groups. (A) Number of parvalbumin-expressing cells in the dorsal hippocampus overall and (B) in hippocampal subfields CA1, CA3 and dentate gyrus across Control, Suc and CafSuc groups. (C) Dorsal hippocampal regions studied, adapted from Paxinos and Watson (2013). (D) Photomicrographs show representative parvalbumin stained neurons in the dorsal hippocampus of Control, Suc and CafSuc treatment groups, 10× magnification, scale bar = 500 μm; inset shows 40× magnification, scale bar = 25 μm. HPC = hippocampus; DG = dentate gyrus; #PV+ cells = number of parvalbumin-expressing cells; Suc = Sucrose group; CafSuc = Sucrose plus caffeine group. \**p* < .05. \*\**p* < .01. \*\*\**p* < .001. Error bars represent +*SEM*.

#### 3.6.2. Basolateral amygdala

Groups showed similar counts of BLA PV+ cells (see Figure 6A; no significant main effect of diet, *F* < 1).

**Figure 6.**
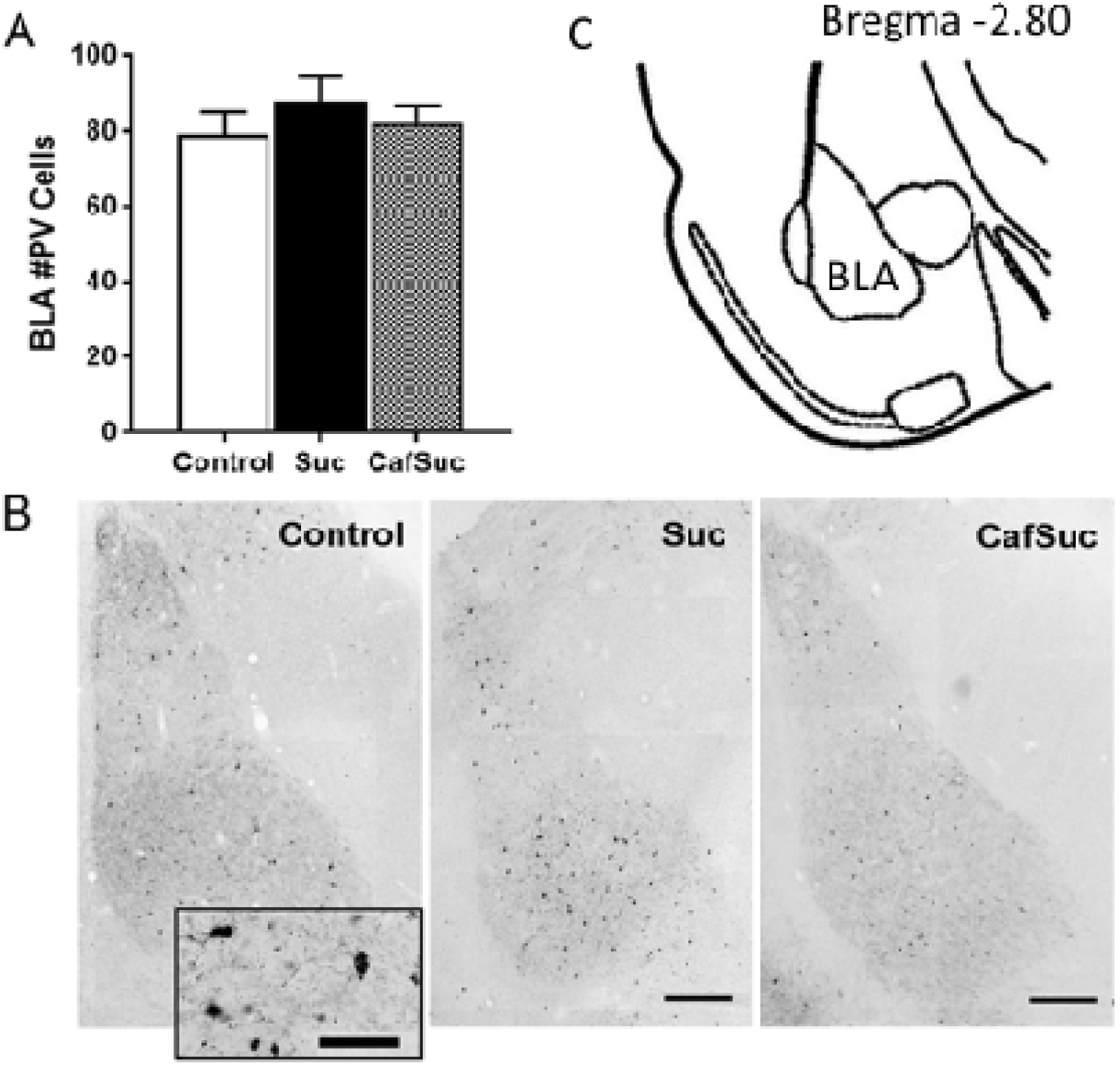
Basolateral amygdala (BLA) parvalbumin immunoreactivity. (A) Number of parvalbumin-expressing interneurons in the BLA across Control, Suc and CafSuc treatment groups. (B) Photomicrographs show representative parvalbumin stained neurons in the basolateral amygdala of Control, Suc and CafSuc treatment groups, 10× magnification, scale bar = 250 μm; 40× magnification, scale bar = 25 μm. (C) Basolateral amygdala (BLA) area studied, adapted from Paxinos and Watson (2013). #PV cells = number of parvalbumin-immunoreactive cells; Suc = Sucrose group; CafSuc = Sucrose plus caffeine group. Error bars represent +*SEM*.

### 3.7. Doublecortin immunofluorescence

Groups differed in %Area_DCX_ (Figure 6), demonstrated by a significant main effect of diet, *F*(2,33) = 8.9, *p* = .001. Relative to Controls, %Area_DCX_ was significantly reduced in CafSuc treatment (*p* = .001), indicating reduced doublecortin immunofluorescence; but %Area_DCX_ did not differ compared to controls in the Suc treatment group (*p* = .17).

**Figure 7.**
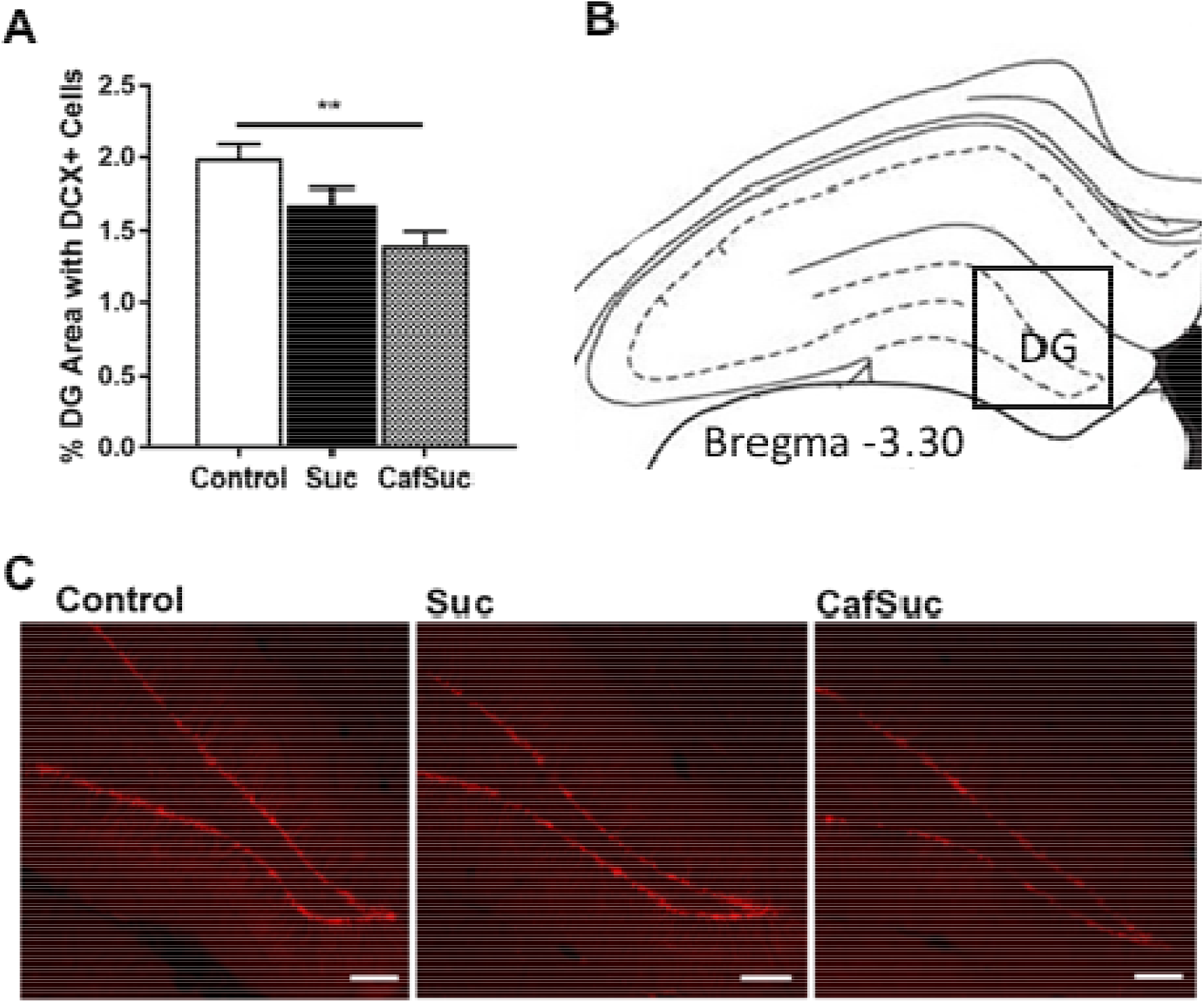
(A) Percentage of area covered by doublecortin-immunofluorescent cells in the dorsal dentate gyrus of the hippocampus across Control, Suc and CafSuc groups. (B) Representation of the 1mm^2^ area of the dentate gyrus measured (at bregrma −3.30mm). (C) Photomicrographs show representative doublecortin stained neurons in the dentate gyrus of Control, Suc and CafSuc treatment groups, 20× magnification, scale bar = 100 μm at approximately bregma −3.30mm. DG = dentate gyrus; DCX+ cells = doublecortin-immunopositive cells. \*\**p* < .01. Error bars represent +*SEM*.

## 4. Discussion

In this study, we examined the effects of intermittent consumption of 10% sucrose or 10% sucrose plus 0.04% caffeine in young male rats on memory, anxiety-like behaviour and parvalbumin and doublecortin immunoreactivity in the dorsal hippocampus.

Consumption of sucrose, or sucrose plus caffeine had differential effects on memory performance in rats. Intermittent sucrose consumption impaired performance of place recognition and long-term novel object memory. In contrast, CafSuc and control treatment groups showed intact place recognition and novel object recognition memory. This indicates that the addition of caffeine to sucrose solution may protect against sucrose-evoked cognitive deficits in hippocampal-mediated memory (Nieradko-Iwanicka et al., 2016) and caffeine may improve hippocampal function (Ardais et al., 2014; Sahu et al., 2013). Furthermore, CafSuc treatment improved long-term object memory performance relative to Suc treatment, demonstrating that the addition of caffeine to sucrose may enhance aspects of memory consolidation. However, as a caffeine only control group was not used in this experiment it cannot be concluded whether the general cognitive enhancing effects in this test were due to caffeine alone or caffeine in combination with sucrose.

All groups showed intact short-term object recognition memory, which is typically dependent on perirhinal, rather than hippocampal, function (Bartko et al., 2007). Previous studies have shown that short-term object memory function in rats was not disrupted following either intermittent (Abbott et al., 2016) or continuous sucrose consumption (Beilharz et al., 2016), or *ad-libitum* caffeine access (Abreu et al., 2011) when objects were perceptually different. However, a recent study has indicated that high sucrose diets can disrupt object recognition performance when objects have increased similarity and share multiple features (Buyukata et al., 2018).

Sucrose and sucrose plus caffeine evoked differential effects on anxiety-like behaviour measured in the EPM and open field depending on whether rats have recently accessed the solutions or not. Rats were tested on the EPM prior to their daily sucrose or sucrose plus caffeine access, in this test rats in the CafSuc and Suc treatment conditions showed increased anxiety-like behaviour (i.e., closed-arm occupancy) compared to control rats. Immediately following their daily 2h access to solutions, CafSuc treatment displayed reduced anxiety-like behaviour (i.e., border occupancy) in the open-field compared to Suc and Control treatment groups. This observation contrasts a previous study that showed continuous caffeine intake increased anxiety-like behaviour in rats (Abreu et al., 2011). The differences in anxiety-like behaviour could arise from the intermittent access procedure used in our current study. As such, recent access to CafSuc may have reduced anxiety in the open field, however, further testing should be conducted to examine whether anxiety-like behaviours differed between groups in the EPM immediately following access to sucrose / CafSuc. The increased anxiety observed in both Suc and CafSuc treatment group suggests that regular consumption of these solutions can impact upon aspects of emotional behaviour. This observation complements human studies correlating energy drink intake and reported anxiety in young people (Stasio et al., 2011; Trapp et al., 2014), and that caffeine improved subjective mood in habitual caffeine consumers (Yeomans et al., 2002).

The sucrose group showed decreased hippocampal parvalbumin immunoreactivity, as previously observed in rats consuming high sucrose diets (Reichelt et al., 2015). However, parvalbumin immunoreactivity in the CafSuc group did not differ to the control group. This suggests that the addition of caffeine to sucrose may protect against sucrose-evoked reductions in parvalbumin immunoreactivity. Hypercaloric diet consumption can reduce expression of brain neurotrophic factor (BDNF) in the cortex (Kanoski et al., 2007; Molteni et al., 2002), however caffeine administration increases cortical BDNF (Lao-Peregrin et al., 2017; Moy and McNay, 2013), and thus may prevent diet-induced BDNF changes, preserving cognitive performance in the CafSuc group. Chronic sucrose consumption alters the morphology of neurons, in particular reducing dendritic spine densities and complexity (Klenowski et al., 2016; Shariff et al., 2017). Caffeine can evoke dendritic morphological alterations to neurons, including increasing spine density in neurons (Vila-Luna et al., 2012), thus the addition of caffeine to sucrose may negate sucrose-induced morphological alterations to neurons. Moreover, chronic caffeine intake antagonises hippocampal A_1_ adenosine receptors, blocking their inhibitory control over parvalbumin-expressing interneurons (Yang et al., 2013). Increased parvalbumin interneuron activity through the combination of caffeine with sucrose may prevent sucrose-evoked memory disruption (Ognjanovski et al., 2017), providing a potential mechanism for the preserved memory performance in the CafSuc treatment group. Further research is needed to determine the neural mechanisms underpinning high sugar diet-induced alterations to hippocampal parvalbumin neurons, and other neurobiological mechanisms affected.

No differences in BLA parvalbumin immunoreactivity were observed between groups despite Suc and CafSuc treatment increasing aspects of anxiety-like behaviour. This contrasts studies showing modulation of BLA parvalbumin-expressing interneuron populations increased anxiety-like behaviour (Godavarthi et al., 2014; Urakawa et al., 2013). However, this indicates that changes in pavalbumin neurons may region specific to the hippocampus and prefrontal cortex, as no changes in the BLA have previously also been noted in rats with intermittent access to hypercaloric foods (Baker and Reichelt, 2016). Further mechanistic insights are required to understand how diet and caffeine consumption alters anxiety-like behaviours.

We observed that CafSuc treatment reduced doublecortin immunofluorescence in the DG, a neuroproliferation marker that has also been linked to anxiety (Yun et al., 2016). Doublecortin immunoreactivity in the DG was significantly reduced in CafSuc rats compared to Controls, suggesting that despite reductions in neuroproliferation, the performance of hippocampal-mediated memory tasks were preserved. This complements previous research showing that 7 days caffeine treatment reduced doublecortin-expressing neurons (Wentz and Magavi, 2009), indicating that caffeine may reduce neuroproliferation. Moreover, as CafSuc rats consumed significantly more solution than the Suc treatment group, this may have increased sucrose-evoked alterations in neuroproliferation.

A caffeine supplemented diet has been shown to increase anxiety-like behaviour and dysregulated hypothalamic-pituitary-adrenal-axis stress reactivity (O’Neill et al., 2016). Further studies could examine levels of plasma corticosterone and cortical corticotrophin receptor levels to establish whether CafSuc differentially impacts glucocorticoid signalling pathways compared to Suc. Moreover, the effect of these treatments on doublecortin and parvalbumin expression within the ventral hippocampus should be explored, as this region is critically linked to the expression and regulation of anxiety-like behaviour (Adhikari et al., 2010; Bannerman et al., 2003).

In this experiment, the CafSuc treatment group consumed significantly more solution than Suc rats from week 2 onward. It is hypothesised that the addition of caffeine may have enhances the reinforcing effect of sucrose, particularly as caffeine can promote dopamine signalling (Nall et al., 2016) and synergistically activates dopamine D2-like receptors (Manalo and Medina, 2018). This provides a putative mechanism that also supports studies in humans demonstrating that the addition of caffeine to sugar-sweetened beverages increases consumption (Keast et al., 2015). Body weight differences were not observed between treatment groups. This may be due to a general reduction in chow consumption leading to the titration of overall energy intake (Kendig et al., 2013). Moreover, the rapid growth occurring during the adolescent period has been shown to be protective against hypercaloric diet-induced weight gain, and has been observed in both rats (Baker and Reichelt, 2016) and mice (Labouesse et al., 2013).

### 4.1. Conclusions and Implications

In this study we provide evidence comparing the behavioural and neurobiological impact of the consumption of sucrose solution and sucrose plus caffeine solution in rats to model the consumption of soft drinks and energy drinks respectively. Collectively, these findings suggest that intermittent access to sucrose solution or sucrose plus caffeine solution increases anxiety-like behaviours prior to expected access. However, following access to solutions no differences between groups were observed in anxiety-like behaviour.

The current study did not have a caffeine only treatment group. Such group was omitted as the presentation of caffeine alone, due to its bitter taste, deterred consumption during the 2 h binge period. Future studies could utilise non-caloric sweeteners, e.g. saccharin, to enhance palatability of the solution, however, saccharin consumption impacts on motivated behaviour in rats (Vendruscolo et al., 2010), and this group may show a different behavioural phenotype.

Studies have highlighted adolescence as a developmental period of vulnerability to hypercaloric diet-induced behavioural changes (Baker et al., 2017; Noble and Kanoski, 2016). As such, adult rats may not show as pronounced neurobiological and emotional changes to Suc or CafSuc treatments. The enduring nature of these effects warrant further investigation, particularly whether the addition of caffeine can restore behavioural changes after extended high sucrose diet consumption.

These data indicate that caffeine may protect against high sucrose diet evoked cognitive impairment, as CafSuc treated rats did not show memory deficits and had comparable parvalbumin immunoreactivity compared to controls. However, CafSuc treatment significantly reduced doublecortin immunofluorescence. These data suggest that regular energy drink consumption potentially alters emotional behaviours and highlights the need for recommended intake guidelines.

## 5. Acknowledgments

ACR receives funding from an Australian Research Council Discovery Early Career Research Award (DE140101071).

## List of abbreviations

ANOVA: Analysis of Variances
BDNF: Brain derived neurotrophic factor
CafSuc: Sucrose plus caffeine
DG: Dentate Gyrus
EPM: Elevated Plus Maze
eR: Exploration Ratio
NHS: Normal Horse Serum
NOR: Novel Object Recognition
NPR: Novel Place Recognition
NRS: Normal Rabbit Serum
PB: Phosphate Buffer
PBS: Phosphate Buffered Saline
PBS-T: Phosphate Buffered Saline plus 0.1% Triton-X
PFA: Paraformaldehyde
PV: Parvalbumin
Suc: Sucrose

